# A measure of agreement across numerous conditions: Reproducibility of co-expression networks across tissues

**DOI:** 10.1101/125450

**Authors:** Alejandro Cáceres, Juan R. González

**Author notes:** Corresponding Author: Alejandro Cáceres,.

## Abstract

There is great interest to study how co-expression gene networks change across tissues. However, the reproducibility assessment of these studies is challenged by a lack of fully confirmatory experiments from independent researchers. While an increment in the number of studies with expression data for several tissues is expected, statistical measures are still needed to assess the reproducibility between studies. We identified a gap in the statistical literature concerning the assessment of agreement between studies across numerous conditions. The gap precluded us to test, using standard statistics, the level of agreement between the GTEX (RNAseq) and BRAINEAC (microarray) studies to distinguish the structure of co-expression networks across four brain tissues. We propose a generalization of a classical measure of agreement, Cohen’s *κ*, derive its distributional characteristics and determine its reliability properties. In the gene expression studies, our generalization of *κ* showed full agreement for genome-wide networks in BRAINEAC benchmarked against GTEX, and highest agreement for brain specific pathways. Our highly interpretable measure can contribute to anticipated efforts on reproducibility research.

## Introduction

Reproducibility is a pressing issue in biomedical research that particularly worries a large number of researchers in the field (Baker, 2016). Research guidelines from leading journals and the American Statistician Association urge for the need of confirmation studies and accurate statistical reporting (www.nih.gov/about/reporting-preclinical-research.htm) (Wasserstein and Lazar, 2016; Mogil and Macleod, 2017). In systems biology, interaction networks are often derived from the integration of high-throughput data and confirmatory studies may be available for simple experimental designs, in public repositories such as GEO (www.ncbi.nlm.nih.gov/geo/). A number of metrics exist to test the preservation of a network under different conditions (Langfelder *et al*, 2011). If the conditions are different experiments then the measures can be used to assess the reproducibility of the network. However, in more complex studies, the condition levels may change within a single study, such as those that aim to identify the structure of a network in different tissues. Clearly, the reproducibility and validity of such observations also need to be assessed. While preservation metrics can be used again as pair-wise comparisons of one network between two experiments, on a single tissue, the overall reproducibility should assess the degree of agreement to identify different network structures across all tissues.

In statistics, there are numerous ways to measure the reliability of an observation. Reliable observations are reproducible and accurate. Agreement measures between two experiments are used to assess the consistency of the observations being made. If observations are classifications of individuals into groups, Coehn’s *κ* and its generalizations are typically used (Cohen, 1960; Banerjee *et al*, 1999); if observations are continuous then a number of correlation measures can be used, such as intra-class correlations (Shrout and Fleiss, 1979). These and other agreement measures are suitable when experiments are performed under comparable or controlled conditions. However, numerous studies are designed to test a group of individuals under a range of varying condition levels. In these cases, it is of interest to assess the reliability of the measures across each of such levels. Remarkably, for this type experiments, there is a lack of agreement measures that, in particular, can help us assess the reproducibility to distinguish the structure of a co-expression gene network across numerous tissues. We, therefore, propose a generalization of Cohen’s *κ* to measure the overall agreement between experiments across a range of conditions.

The GTEX project is an unprecedented effort to study the gene expression in tens of tissues in hundreds of subjects (Lonsdale *et al*, 2013). It is therefore a strong candidate for becoming a preferred benchmark for interaction networks inferred in different tissues. Currently, the validity of a gene or protein network derived from high-throughput data is often benchmarked against networks derived from current knowledge of specific interactions, given by curated pathways, specific experiments or even text mining of published articles (Szklarczyk *et al*, 2014). This type of confirmatory analysis extract networks that are a mixture of interactions individually reported on different tissues. Therefore, while validity is investigated, in terms of consistency with previous knowledge, reproducibility on a given tissue is not being measured. Reproducible networks are observables of reproducible experiments. As the number of studies with expression data in multiple tissues is expected to increase, agreement measures against GTEX may serve as a reproducibility assessment of network structure across tissues (Mogil and Macleod, 2017).

Some studies that measure gene expression in different tissues are already available, one of which is the BRAINEAC project (Trabzuni *et al*, 2011). Here, the gene expression using microarray data was measured in hundreds on un-demented individuals at the time of death in nine different brain tissues. Using our agreement measure, we therefore investigated the reliability, between BRAINEAC and GTEX, of discriminating gene networks across four brain tissues. We tested the reliability of genome-wide gene network and 287 KEGG pathways (www.genome.jp/kegg).

## Results

We propose a measure of agreement between two studies to discriminate observations (network structures) across numerous conditions (tissues). Fig 1 illustrates a simulated example where a reference experiment with three condition levels is tested for reproducibility with one successful and one failed efforts. For experiments with successful reproducibility, we expect network correlations between experiments to be maxima when networks are inferred on the same tissue. Failed reproducibility should show network correlations on the same tissue not different than any other. A proposed measure of overall reproducibility, λ computed for this example (see methods), is an estimator of the probability that network correlations between experiments on the same tissue are maxima.

**Figure 1.**
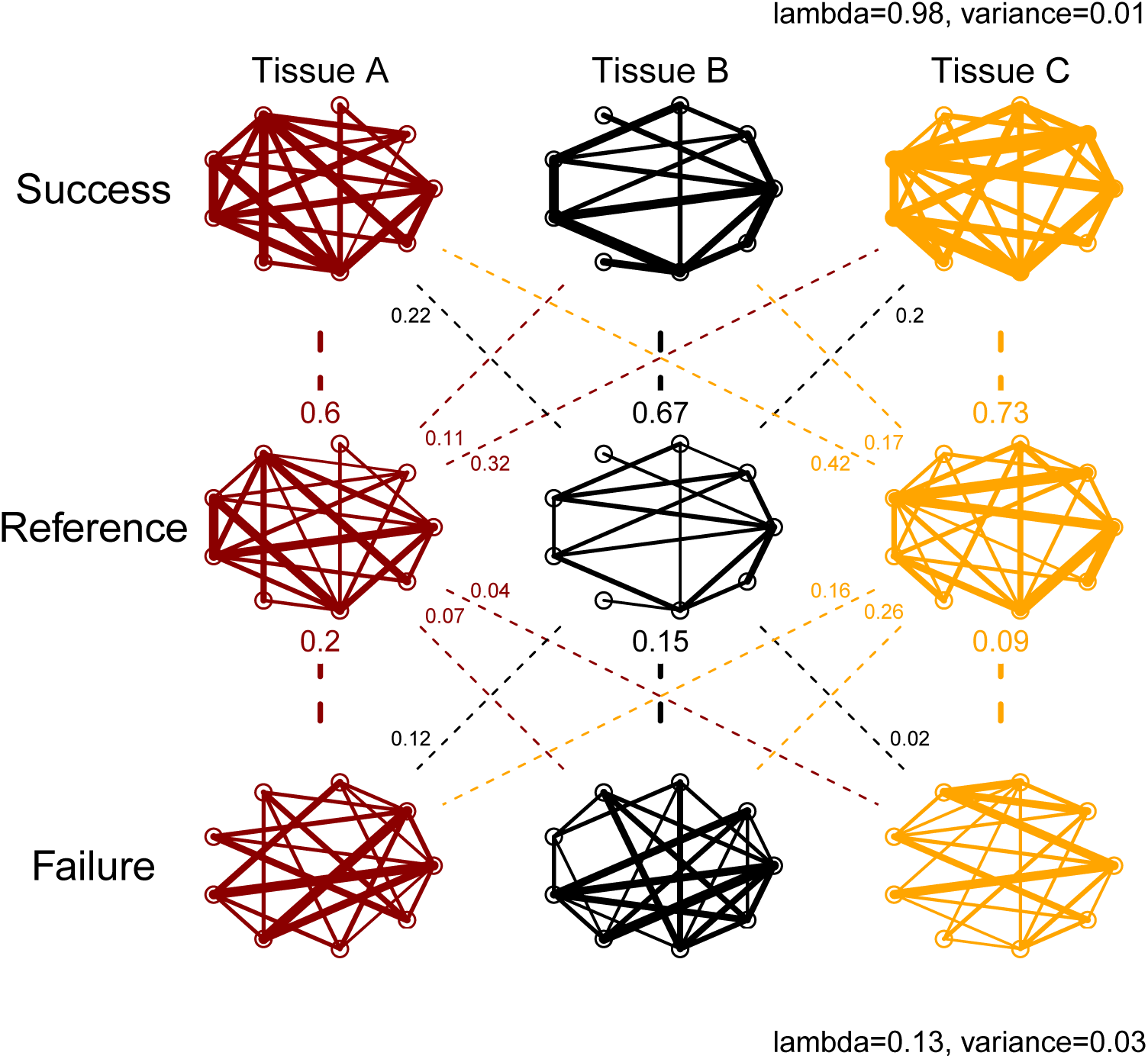
Reproducibility measurement between simulated studies: Reference, Success and Failure. Gene coexpression networks of three tissues (A, B and C) and 9 genes are inferred in each study, which may be based on different population samples and experimental setups. Solid lines represent gene-pair correlations in the network, where line thickness is scaled by correlation strength. Between studies correlations are represented by broken lines. λ is a measure of overall agreement between two studies: Reference Vs Success (top) and Reference Vs Failure (bottom).

### Properties of λ through simulations

Suitable reliability measures satisfy three basic properties: i) their values range from 0 to 1; ii) they tend to 0 when no agreement is expected and tend to 1 when full agreement is expected and iii) they account for random agreement. We studied the properties of λ with extensive simulations. While λ is applicable to more general situations than those covered by *κ*, we compared the performance of λ with Cohen’s *κ* in simulated cases where both measures can be used.

We recreated reproducibility assessment for three different number of tissues *k* = (5, 10, 15) and three possible forms for the cross-tabulation of agreement between experiments (methods, expressions 4 and 5): marginal equiprobability 1/*k* for all tissues (scenario 1), marginal probability of ∼ 1/*j* for tissue *j* (scenario 2), and marginal probability ∼ 1/*j*^2^ (scenario 3), where *j* = 1, …*k*. Scenarios 2 and 3 were designed to test situations where high correlations in the case of λ, or measurements on the case of *κ*, tend to concentrate around one tissue *j* = 1. We performed 10,000 simulations under different initial conditions that allowed us to cover the full range expected agreement (*P*_0_, in methods) from 0 to 1.

We first confirmed that λ ranged from 0 to 1, as it is the expected value of a probability, holding basic property i). We then observed that when null agreement was expected (*P*_0_ = 0) λ tends to 0, while it tends to 1 for full agreement (*P*_0_ = 1), as required by property ii). Consitently with this, we found that λ increased monotonically with *κ* for all the simulation scenarios, see Fig EV1. The functional dependence was highly stable under different scenarios, revealing, as expected, high λ agreement for fair values of *κ* (0.2, 0.4), given that the latter is a measure of exact agreement rather than discriminative agreement. Therefore, agreement as measured by λ has more power than agreement measured by *κ*.

For low values, λ tends to zero when *κ* can take small negative values, a situation already described in Cohen’s work (Cohen, 1960). We also observed that for a given *κ* there is a sizable range of λ values, in particular as tissues become less equiprobable (scenario 3). We noted that if the number of tissues is small (*k* = 5) and the marginal distribution greatly concentrates around one single tissue (*j* = 1 for scenario 3), then λ tends to 1/*k* (0.2) because the experiments can clearly distinguish that tissue from the rest. In this case, *κ* tends to zero.

In relation to basic property iii), we studied the relationship between the agreement measures that account for random agreement with those that do not. In our simulations, we confirmed that *κ* is lower than *P*_0_ (see Fig 2); which illustrates the initial motivation for *κ*’s definition of a measure that corrects for random agreement (Cohen, 1960). Similarly *r*, the fraction of times the diagonal terms in (methods, 4) are row and column maxima, is higher than λ, a distributional estimate of such fraction. Note also that the range of *r* is discontinuous with *k* + 1 possible values, while λ is a continuous value from 0 to 1.

**Figure 2.**
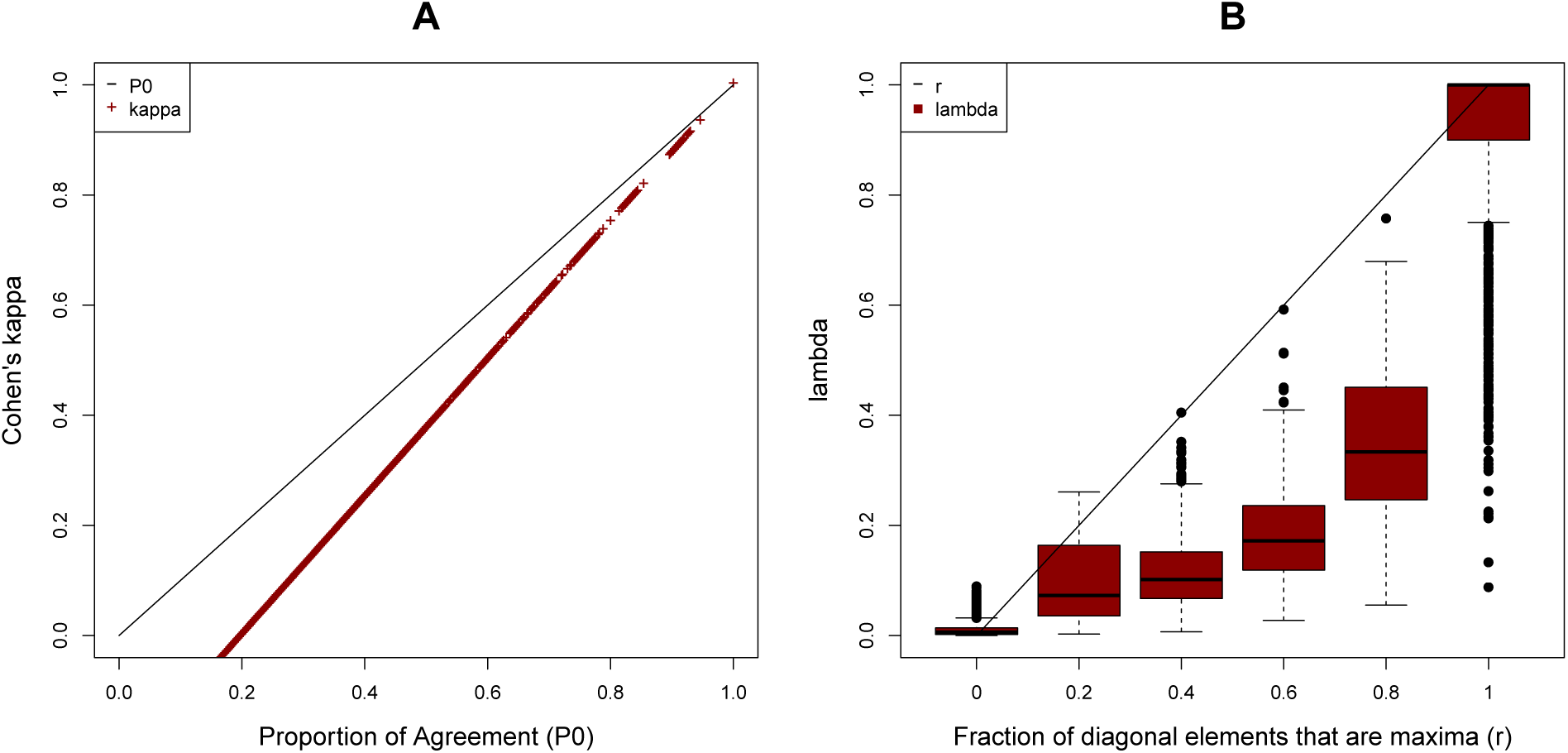
Comparison between measures that correct for random agreement with those that do not. A Cohen’s *κ* is compared with *P*_0_ (the total proportion of agreement). B λ is compared with *r* (the total fraction of times the diagonal elements in the cross-tabulated table of correlations are row and column maxima).

We finally studied λ’s variance and found that it decreases with the number of tissues, and departure from marginal equiprobability (Fig EV2). We observed that while *κ* has a one-to-one correspondence between mean and variance, for a given λ a range of variances are allowed; see Fig EV3. In particular, *κ*’s variance are minimum at extreme values (0,1). λ’s variance, in contrast, decreases towards zero for a range of values.This occurs when the mean of λ tends to *r*, that is, when the probabilities of diagonal terms in (methods, 4) of being maxima tend either to zero or to one. From a practical point of view, this means that if the elements of the cross-tabulated table of correlations between experiments (4) are determined each with high precision (low variance) then the agreement measure can also be estimated with high precision. The effect is clearly visible in the scenario 2 and low number of tissues. As the number of conditions increase, the effect should be visible with a substantial increase on the number of simulations. When concentration of marginal probability around a single tissue is present, we observed a clear reduction of the possible values for the variance around λ = 1/*k*.

### Genome-wide gene network

We inferred the genome-wide co-expression network for 10,683 genes across the GTEX and BRAINEAC studies in four brain tissues: cerebellum, frontal cortex, hippocampus and putamen. The network was fully characterized by 5.7 × 10^7^ interactions which correspond to the elements of the upper triangular correlation matrix between expression levels. We assessed the agreement between studies to distinguish the structure of the genome-wide network across all four tissues. Fig 3 illustrates the correlations between studies across tissues. We observed that all correlations were similar in size between (0.37, 0.46). However, their standard errors were small (∼ 10^−5^), given the large number of degrees of freedom. More specifically, the figure shows that the cerebellum and frontal cortex diagonals are the maxima of their rows and columns, and therefore the two studies can discriminate between them. For the hippocampus and putamen, note that they are the second maxima after their correlations with functional cortex in GTEX. Therefore, the experiments cannon clearly agree on how distinguishable the network is between the frontal cortex, the hippocampus and putamen.

**Figure 3.**
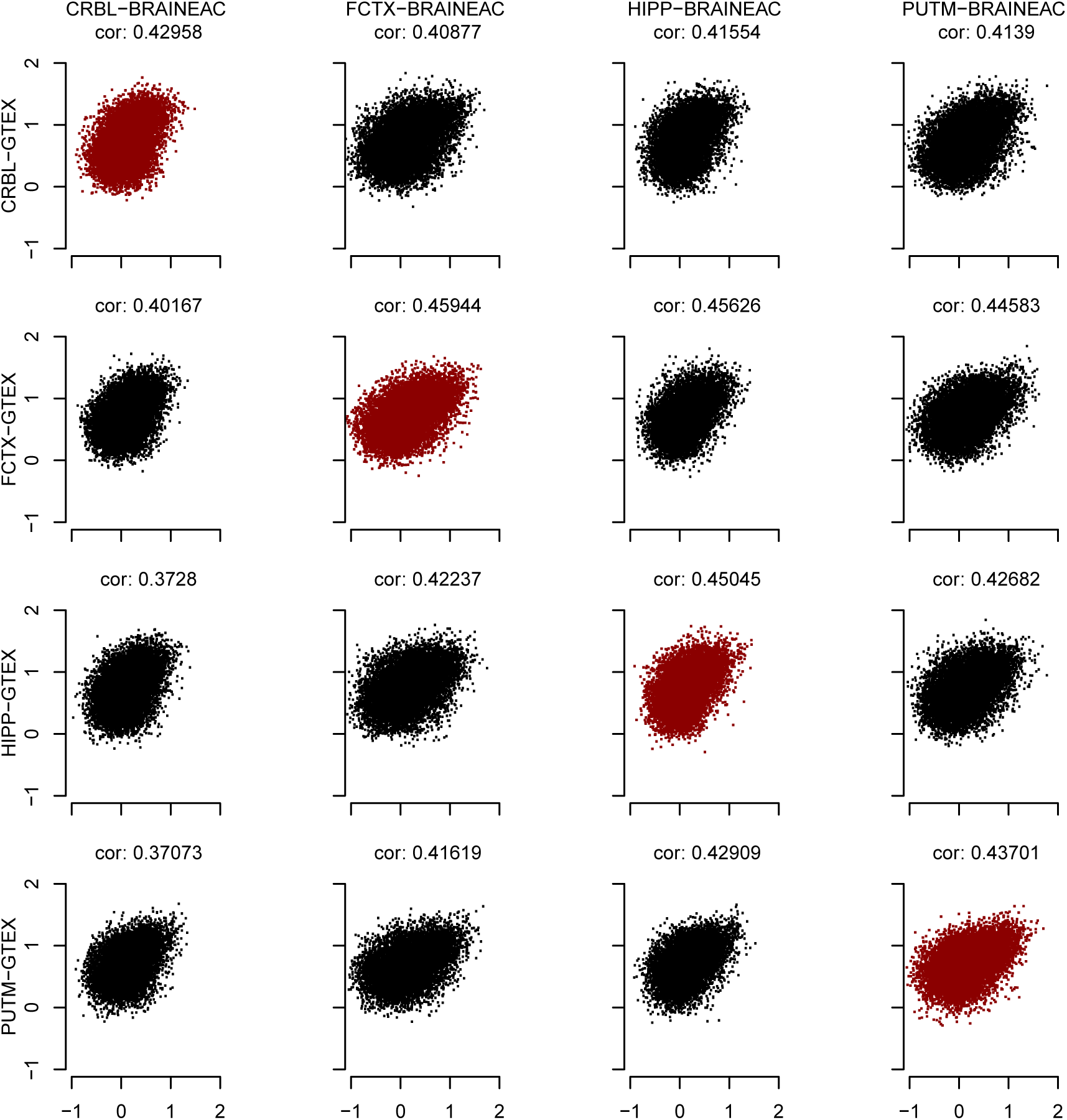
Correlation matrix between the GTEX and BRAINEAC studies across four brain tissues (CRBL:cerebellum, FCTX:frontal cortex, HIPP: hippocampus, PUTM: putamen). The diagonal terms are shown in red. The agreement measure λ assesses the expected fraction of times the diagonal terms are row and column maxima, given the distribution of the correlation estimates.

We computed the agreement measure λ from normally distributed estimates derived from the between study correlations (Appendix Tables S1 and S2). As expected from the observations made in Fig 3, we obtained λ = 0.5 with vanishing variance. This value of λ reports that a fraction of 2/4 conditions are agreed to be different between experiments. The high precision of the estimate follows from the small standard errors of the correlations, due to the large number of degrees of freedom.

We also benchmarked BRAINEAC networks with respect to GTEX. We hence assessed if the diagonal terms were maxima within their rows only (see methods). In this case, we confirmed that all diagonal terms were their row maxima (see Table S3), and therefore λ = 1. These result show that, leaving other confirmatory studies to establish GTEX as a possible benchmark, BRAINEAC fully agrees with GTEX in terms of sensitivity and specificity.

### KEGG pathways

The Kyoto encyclopedia of genes and genomes (KEGG) offers a list of experimentally characterized bio-chemichal pathways. We selected the annotated genes in each study, for the proteins of 287 pathways. For the pathways, with more than 5 genes, we computed the full agreement measure λ and its benchmark version,similarly to the previous section.

The full agreement λ is shown in Table 1 that includes pathways with the top values (λ > 0.5). We observe 8 pathways (2%) with agreement between (0.5, 0.75); those are pathways for which there is agreement to discriminate network structures across two and three tissues. No pathway is likely to be reliably different in all four tissues. Interestingly, 5 of these pathways are directly linked with signaling processes specific to brain. The top hit with λ = 0.68 and 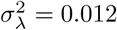, *neuroactive ligand receptor interaction*, is illustrated in Fig EV4. We observed that the cerebellum is not clearly distinguishable by BRAINEAC, as the diagonal term is the minimum in the row. However, a clear distinction is obtained for the frontal cortex, hippocampus and putamen areas. The estimate for λ was lower than *r* = 0.75, as it accounts for sizable uncertainty in the estimates of the correlations.

**Table 1:** Agreement measure λ between BRAINEAC and GTEX for distinguishability of KEGG pathways in more than 2 brain tissues out of 4 (lambda > 2/4 = 0.5)

| lambda | variance | Description | Ref |
| --- | --- | --- | --- |
| 0.682 | 0.012 | Neuroactive ligand-receptor interaction | hsa04080 |
| 0.655 | 0.024 | Nicotine addiction | hsa05033 |
| 0.600 | 0.046 | Long-term potentiation | hsa04720 |
| 0.579 | 0.015 | Calcium signaling pathway | hsa04020 |
| 0.560 | 0.032 | GnRH signaling pathway | hsa04912 |
| 0.543 | 0.038 | MicroRNAs in cancer | hsa05206 |
| 0.539 | 0.038 | Alcoholism | hsa05034 |
| 0.501 | 0.035 | Transcriptional misregulation in cancer | hsa05202 |

We also benchmarked BRAINEAC with respect to GTEX for the KEGG pathways. We confirmed the higher estimated of λ in this case, since lower comparisons for the diagonal terms are included, and therefore their probabilities of being row maxima increase. In particular, we observed that 5 pathways had agreement between (0.75, 1), meaning that BRAINEAC can agree to distinguish between 3 to 4 tissues in these pathways, if GTEX variability is not taken into account (Table S3). Three of the four pathways are specific to brain and were previously obtained in the full agreement measure. In particular, *neuroactive ligand receptor interaction*, increased to λ = 0.805. We interpret this result as a gained distinction between the frontal cortex, hippocampus and putamen, and an increment in the probability that cerebellum is a maxima; respect to the full agreement measure.

## Discussion

We propose a new measure, λ, of agreement between studies. The motivation of the measure is the assessment of agreement between studies that test the effects varying condition levels on a set of items (subjects, coexpression pairs, etc). We showed through simulations that the statistics conformed to agreement measure properties and we compared it with Cohen’s *κ*. However, formal proves and properties of the statistics are still needed to gain more insight into the measure. Here, we illustrated the large potential of λ applicability,as it carries all the interpretability of *κ* to more general studies.

We specifically studied the agreement between studies to discriminate co-expression network structures across four different brain tissues. We are unaware of similar measures of agreement, in particular, for testing the structure of a network across tissues. Measures of module network preservation allow the reliability assessment of the network over studies, or the preservation of the network between tissues (Langfelder *et al*, 2011). Here, we were interested in assessing the structure of a network between two conditions: studies and tissues; that is, the reproducibility of the network structure across tissue. As the new measure is conceptually closer to inter-observer agreement measures, we designed a simulation framework for which the properties of λ could be compared with those of Cohen’s *κ*. We observed that λ is a suitable reliability measure and, as compared with *κ*, λ systematically leads to higher agreement. Perfect agreement for *κ* is exclusively given by diagonal tables, while perfect agreement for λ is given by maxima diagonal terms in tables where the terms, irrespective of their magnitude, are estimated with sufficient low variance. This is an important difference between the measures, which allow λ to be utilized in more general situations where the elements of the cross-tabulated table between studies are inferences, and not only the proportion of times two raters agree on a measurement of a set of items. In particular, we observed that λ can be estimated with low variance for intermediate values of agreement, or intermediate fraction of the number of tissues, that are reliably different between studies. Therefore, while λ can be less conservative measure, it allows for a suitable generalization to studies that deal with numerous condition levels (tissues); a situation that cannot be assessed with *κ*

In our application to co-expression networks in brain, we found that GTEX and BRAINEAC agree on the discrimination of 2 tissues out of 4, for the genome-wide network. Note that the two studies are based on very different technologies (RNA-seq and microarray) and analysis methods to infer the networks in two different sets of subjects. Our results therefore fully test inter-observer reproducibility. Previous work have tested these two technologies on same subject sample to assess the level of agreement between gene expression measurements (Trost *et al*, 2015; Guo *et al*, 2013). These are important studies to validate experimental techniques. Testing inter-observer reproducibility of network inferences, however, further requires confirmation from from independent experiments on different population samples.

We made two further observations. If GTEX is considered as a benchmark study, the agreement measures increased. In this case, we assume that GTEX validity as benchmark for gene-network inferences should be evaluated in other studies, involving futher experimental validation. We also observed that multiple biochemical pathways can also be assessed for agreement. This focused approach lead to the identification of pathways specific to brain’s biology. Our results suggest that agreement assessment can also be used to identify biochemical pathways with interesting structures that reliably change across tissues.

## Materials and methods

We propose an agreement measure of experiments to discriminate condition levels. While the measure can be applied on different research settings, such as follow-up studies, we illustrate how its need arises from an example in current functional genomic research.

### The problem

Let us assume, without any loss in generality, that we have two experiments that measure genome-wide gene expression in two different population samples, in the same range of tissues, using different experimental setups. For instance, one experiment may use RNA-seq and the other a microarray technology, together with other uncotrolled experimental conditions such as batch effects. We are interested in inferring the co-expression structures of a gene network across tissues and determine whether they are consistent between experiments. The co-expression between two genes in the network is given by their correlation over the subjects’ gene expression levels. Figure 1 illustrates the situation, where co-expression between 9 genes (variables) is shown for 3 tissues (condition levels) in three different studies. Our aim is then to propose a measure of the overall reproducibility of the network inferred between two experiments across tissues.

A gene network can be represented by a correlation matrix between all gene-pairs. Given that the correlation matrix is symmetrical, the network is fully determined by the upper triangular terms of the matrix. Let us assume that for tissue A the gene-pair elements that determine the co-expression network (*netA*) are given by

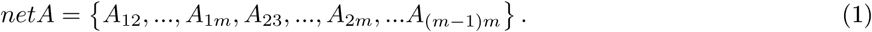
where *m* is the number of genes in the network and the total number of gene-pairs in *netA* is number of elements in the upper triangular correlation matrix 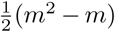. In another experiment, the same gene-pairs correlations are computed

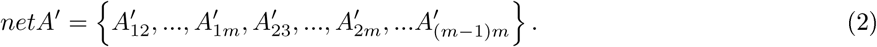
One measure of module preservation is the correlation of the networks between experiments; other presernation measures are also possible (Langfelder *et al*, 2011). We therefore compute the correlation

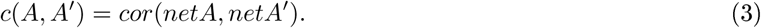
as a measure for the preservation of the network between experiments in tissue A. To assess the overall reproducibility of the network between experiments across all tissues (*A, B* and *C*), we then form the cross-tabulated table of correlations:

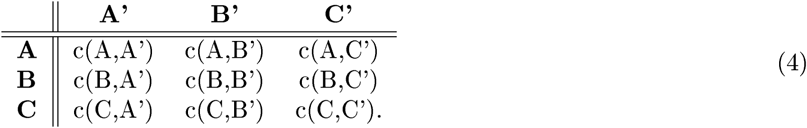
We would then like to have a measure of the agreement from the cross-tabulation (4), whose elements are point estimates with given distributional properties.

### A solution

Cross-tabulation for two judges observing *l* items in *k* categories (*A, B* and *C*) takes a similar form of expression (4),

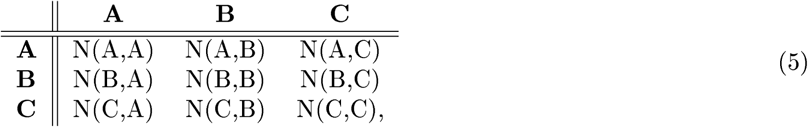
where *N*(*X, Y*) is the number of items measured in category *X* and *Y* by the first and second judge respectively, and ∑_*X,Y*_ *N*(*X, Y*) = *l*. Agreement is typically measured by Cohen’s *κ*

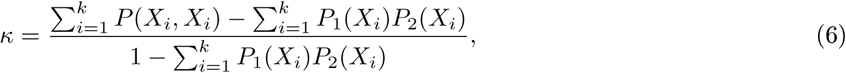
where *P*(*X_i_, X_i_*) = *N*(*X_i_, X_i_*)/*k* is the observed frequency of items that where measured in category *X*_*i*_ by both judges and *P_d_*(*X_i_*) is the frequency of items in *X_i_* observed by judge *d* (*d* = 1, 2). *κ* measures the fraction of discordant observations expected by chance that are actually observed in agreement. The sum *P*_0_ = ∑_*i*_ *P*(*X_i_, X_i_*) is the total fraction of agreement: the proportion of observations that falls in the diagonal and does not account for random agreement.

From the cross-tabulation 4, we propose to measure the probability that the diagonal items on the table are their row and column maxima:

- *p_AA_* = *Pr*(*c*(*A, A′*) > *c*(*A, B′*), *c*(*A, C′*), *c*(*B, A′*), *c*(*C, A′*)),
- *p_BB_* = *Pr*(*c*(*B, B′*) > *c*(*B, A′*), *c*(*B, C′*), *c*(*A, B′*), *c*(*C, B′*)) and
- *p_CC_* = *Pr*(*c*(*C, C′*) > *c*(*C, A′*), *c*(*C, B′*), *c*(*A, C′*), *c*(*B, C′*)),
 where *p_ii_* (*i* = *A, B, C*) is the probability that the correlation in tissue *i* between experiments is the maximum amongst the correlations between the network in *i*, in one experiment, and any other tissue, in the other experiment. These probabilities can be computed as the product of the individual pair-wise probabilities

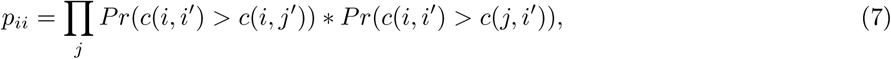
Where the first factor is the maximum over rows (experiment 1), the second factor is the maximum over columns (experiment 2), and the product runs over all other possible tissues (*j*). If we assume that the correlations *c*(*i, j′*) can be transformed to normal random variables *z_ij′_* using, for example, a Fisher’s z transformation, then the probability that the diagonal term (*i, i′*) is higher than other term *j′* in the row can be computed from

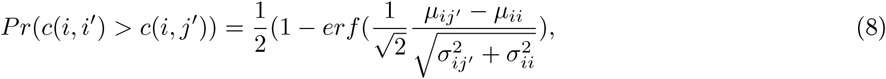
where *erf* is the error function. The expression follows from assuming a transformation *T* such that

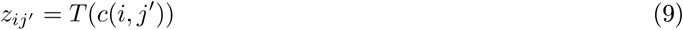

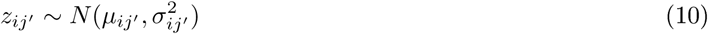
and performing the integration over the joint distribution

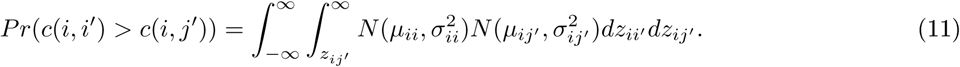
Therefore, we have that the probability that the diagonal term *c*(*i, i′*) is the maximum in the row *i* is

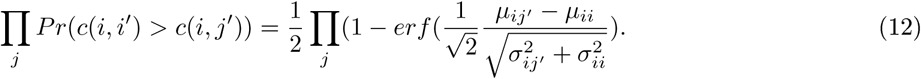
The the probability that the diagonal term *c*(*i, i′*) is the maximum in the column *i* follows a similar form.

Our agreement measure then follows from the overall probability that the diagonal items on the cross-tabulated table are their row and column maxima. This is the probability of *k* successes in *k* Bernoulli trials each of which has its own probability *p_ii_*, or a binomial Poisson distribution with mean and variance

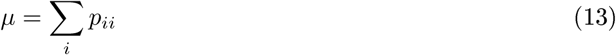

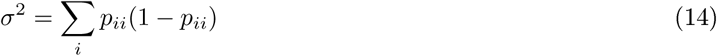

We define the fraction of successes λ = *μ/k* with corresponding variance 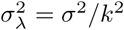 as the agreement measure between experiments to discriminate network structures across conditions. In the case that experiment 1 (in rows) is the benchmark for experiment 2 (in columns), then one is interested in testing whether the diagonal terms are the maxima of their rows only, generalizing the concepts of sensitivity and specificity for more than two conditions. In this case λ can be computed by simply setting *Pr*(*c*(*i, i′*) > *c*(*j, i′*) = 1. Note also that it is straightforward to generalize the measure for more than two experiments by expanding the products in equation (7).

### Comparison between measures of agreement

While λ is a generalization of *κ*, to be applied in more general cases, we compared them in a case where both measured can be computed. Note that obtaining a cross-tabulation (5) from (4) is not univocal. However, a cross-tabulation of (4), where λ is computed, can be obtained from the cross-tabulation (5), where *κ* is defined. Given that for row *i* in (5) the number of observed items is *N_i_* = *P*_1_(*X_i_*)*k*, we can then assume that *N*(*X_i_, X_j_*) is one draw of a binomial process

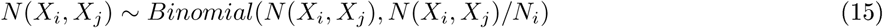
with mean, and variance of the mean,

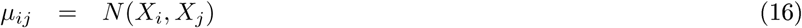

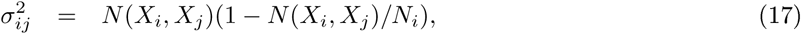
which distributes normally for large *N_i_*. These values can be used in equation 12. With a similar computation for the column elements, the measure λ can be obtained for a table in the form (5) and compared with the value of *κ* for varying values of the total fraction of agreement *P*_0_.

As *κ* is an agreement measure that corrects *P*_0_ for random agreement, we compared λ with the uncorrected agreement measure *r* defined as

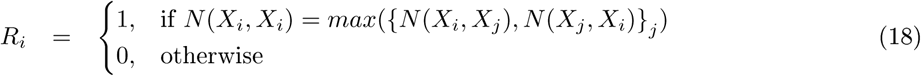

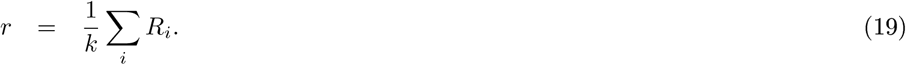
that measures the fraction of times the diagonal elements are thier row and column maxima.

We performed a series of simulations to study the properties of λ with respect to *κ* and *r*. Simulations were obtained for three possible number of condition levels *k* = (5, 10, 15), and three possible forms for the marginal frequencies *P*1(*X_j_*) and *P*2(*X_j_*)

- Senario 1 (equiprobable): 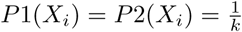, ∀_*i*_
- Senario 2: 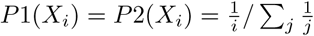
- Senario 3 (the least equiprobable): 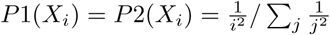

We set the number of observations to *l* = 500. For each scenario, we simulated 50 cases of perfect agreement tables (*P*_0_ = 1), i.e. diagonal matrices, and 50 cases of perfect disagreement (*P*_0_ = 0); those are tables with zeros on the diagonal terms except for the cell of maximum joint probability. For each case, we permuted 20 pairs of observations 100 times, such that the original marginal frequencies were conserved. After each 20 pairs of permutations, we computed the four agreement measures. This procedure allowed the assessment of 10,000 simulations, in each scenario and tissue level, covering the whole agreement interval of *P*_0_.

We used `R. 3. 30` and the package `psych` to perform calculations and compute the Cohen’s *κ*.

### Gene expression data

We downloaded expression data from the GTEX project obtained from RNA-seq (http://www.gtexportal.org). Reads per kilobase per million mapped reads (RPKM) of version 6 were obtained for all brain tissues (`GTEx_Analysis_v6_RNA-seq_RNA-SeQCv1.1.8_gene_rpkm.gct.gz`). Covariates for each tissue were also downloaded (`GTEx_Analysis_V6_eQTLInputFiles_covariates.tar.gz`).

We also downloaded the brain expression data of the BRAINEAC project (http://www.braineac.org/) obtained from winsorized values of exon array data (Affymetrix Human Exon 1.0 ST array). Downloaded data had been previously normalized and corrected for batch effects.

We identified four brain tissues common in both data-sets and for which GTEX had covariate information. Those were cerebellum (CRBL) with 125 individuals in GTEX and 130 in BRAINEAC, frontal cortex (FCTX) with 108 and 135 individuals, (HIPP) hippocampus with 94 and 130 individuals, and putamen (PUTM) with 82 and 135 individuals, respectively. Between the two studies, we mapped 10,683 genes for wich we computed their pair-wise correlations of expression levels. We computed partial correlations for GTEX data, in which we adjusted for the covariates, and a Pearson’s correlation for gene co-expressions in BRAINEAC.

## Acknowledgements

This work was partially supported by Red Española de Supercomputación (BCV-2016-3-0002) and by Ministerio de Economía e Innovación (Spain) (MTM2015-68140-R).

## Author Contributions

AC designed the study, analyzed the data and wrote the manuscript, JRG supervised the work and revised the manuscript.

## Conflict of interest

None declared

**Figure EV1** Comparison between λ and Cohen’s *κ* for values of *P*_0_ (total agreement fraction), ranging from 0 to 1, for varying number of tissues and three different cross-tabulation scenarios.

**Figure EV2** Variance of λ, for 3 tissues and 3 cross-tabulation scenarios, as a function of its mean. The figure illustrates how λ can achieve precise estimates for intermediate agreements.

**Figure EV3** Variance of *κ* and λ as function of their mean values, for scenario 2 and 5 tissues.

**Figure EV4** Correlation matrix between the GTEX and BRAINEAC studies across four brain tissues (CRBL:cerebellum, FCTX:frontal cortex, HIPP: hippocampus, PUTM: putamen) for the neuroactive ligand-receptor interaction pathway. The diagonal terms are shown in red.

